# 3D Printing of Multiscale Scaffolds with Microtopography for Guiding Tissue Organization and Regeneration

**DOI:** 10.1101/2024.09.26.615287

**Authors:** Rao Fu, Evan Jones, Boyuan Sun, Guillermo Ameer, Cheng Sun, Yonghui Ding

**Author notes:** E-mail address (Y. Ding) (C. Sun) (G. Ameer).

## Abstract

Engineering biomaterial scaffolds with multiscale structures, integrating anatomically accurate macroscale architecture (millimeters to centimeters) with microtopographic features (sub-microns to tens of microns), is critical for guiding cellular organization and tissue regeneration. However, fabricating such multiscale scaffolds remains challenging due to the limitations of conventional manufacturing techniques and the trade-off between speed and resolution in current 3D printing methods. Here, we present a multiscale micro-continuous liquid interface production (MµCLIP) technique that enables rapid, one-step 3D printing of centimeter-scale scaffolds with spatially tunable microtopography of various sizes and geometries in just a few minutes (up to 1 mm/min). To showcase the versatility of our technique, we printed a one-centimeter-long tubular scaffold with dual microtopographic patterns, i.e. 20 µm axially aligned grooves on the internal surface and 15 µm circumferentially aligned rings on the external surface. These scaffolds induced the simultaneous orientation of vascular endothelial cells along the axial grooves and vascular smooth muscle cells along the circumferential rings, mimicking the orthogonally aligned bilayer architecture of natural arteries. Moreover, the groove patterns significantly accelerated endothelial cell migration, potentially enhancing endothelialization in vascular implants. This approach provides a versatile tool for designing advanced scaffolds and medical devices that harness microtopography to guide tissue organization and enhance regeneration.

---

The goal of regenerative engineering is to repair or restore the function of diseased or damaged tissues and organs, often accomplished through the use of biomaterial scaffolds. These scaffolds are engineered to provide mechanical support and guide cell behavior, facilitating the formation of neo-tissues that replicate the mechanical and biological functions of their native counterparts. Many tissues in the body, such as arteries, nerves, muscles, myocardium and intestines, exhibit specific spatial orientation of cells and matrices, which are crucial for their mechanical and biological functions^1-3^. For instance, the longitudinal alignment of myofibers in skeletal muscles is integral to supporting movement and enduring mechanical loads^4^. Similarly, in the nervous system, the alignment of Schwann cells is vital for guiding axonal regrowth^5^. Moreover, an orthogonally aligned bilayer architecture is present in the walls of intestines and arteries. In arterial walls, the circumferential alignment of smooth muscle cells (SMCs) in the media layer enables an effective vessel contraction and dilation^6, 7^. While the axial orientation of endothelial cells (ECs) in the intima layer of arterial wall is vital for maintaining vessel homeostasis and their atheroprotection roles^8, 9^. The ability to regenerate tissues in a way that mirrors such spatial orientation is essential for reinstating their unique functional characteristics. However, achieving precise control over the cellular and tissue in three-dimensional (3D) scaffolds, particularly for multiple cell types with distinct orientations, remains a significant challenge.

Various engineering strategies have been developed to control cell alignment, including mechanical loadings^10^, chemical patterns or gradience^11^, and surface topography^12^. Among these, surface topography, especially ordered arrays of patterns at the scale of individual cell size (microns to tens of microns), has proven to be a powerful tool for guiding spatial orientation and function at both the cellular and tissue levels without the need of external stimuli^13^. Nonetheless, most studies on topographic effects have been limited to two-dimensional (2D) planar substrates, primarily due to the difficulty of creating topographic micropatterns on curved surfaces of 3D structures using conventional micro-/nano-fabrication techniques like photolithography^14^. To overcome this challenge, various manufacturing methods have been explored, including short-pulsed laser processing^15^, sacrificial templating^8, 16^, and modified electrospinning ^17, 18^. Although these approaches have demonstrated the ability to guide the biomimetic orientation and functional recovery of tissues such as muscles, nerves, and arteries^16^, they suffer from significant drawbacks, including low fabrication efficiency, limited reproducibility, and the complexity of multi-step processes, which hinder their clinical adoption. For instance, short-pulsed laser processing relies on point-by-point scanning, limiting its efficiency, especially for complex structures with limited internal access^19, 20^. Sacrificial templating and electrospinning involve time-consuming procedures and are often restricted to simple shapes, such as cubes or cylinders. Thus, there remains a need for a simple and effective strategy to produce 3D scaffolds capable of guiding cell and tissue orientation.

3D printing offers substantial flexibility for the fabrication of complex 3D geometries, enabling the production of patient-specific scaffolds for regenerative engineering. However, most existing 3D printing technologies face trade-offs between resolution, building volume, and production speed. Techniques such as two-photon polymerization can achieve submicron resolution but are slow, taking weeks to fabricate centimeter-scale objects^21, 22^. While methods like projection-based stereolithography and continuous liquid interface production (CLIP) have significantly improved fabrication efficiency^23, 24^, producing muti-scale 3D objects with features ranging from the centimeter to the micro-/nano-meter scales remains a formidable challenge. To address this, two main strategies have been explored: ink chemistry-based methods and improvements in manufacturing process or hardware. Ink chemistry-based methods, such as chemically mediated phase separation, have been used to generate micro/nanoscale pores throughout 3D-printed objects^25-27^. However, these strategies are heavily reliant on precise ink formulation and provide limited control over the geometry, size, and spatial distribution of the porous structures. Additionally, producing distinctive micro/nanoscale structures at different spatial locations within the same object remains highly challenging. On the other hand, improvements in manufacturing processes or hardware have aimed to improve resolution while maintaining high fabrication efficiency. Despite significant advancements, most existing multiscale 3D printing methods are limited by achievable feature sizes ranging from hundreds of microns to cetimeters^28, 29^. However, to effectively guide cell alignment and other behaviors, topographic features need to be on the scale of individual cells, typically ranging from submicron to tens of microns^13^. DeSimone’s group has introduced a high-resolution CLIP printer capable of producing multiscale objects at high speed by implementing a reduction lens optic system and a contrast-based focusing system^30^. While they demonstrated the fabrication of aligned micropatterns as small as 7.5 µm, it is limited to creating 2D patterns in the *xy*-plan (i.e., normal to the printing direction).

In this study, we aimed to develop a facile and efficient 3D printing approach to fabricate multiscale scaffolds featuring spatially controlled microtopography for guiding the biomimetic tissue orientation and regeneration. We introduced a multiscale micro-continuous liquid interface production (MµCLIP) method that enables the one-step fabrication of centimeter-scale 3D scaffolds with microtopographic patterns of various sizes (12-80 µm) and geometries (grooves, pillars, and rings) at a high speed (1.0 mm/min). We showcased that 3D-printed tubular scaffolds with dual microtopographic patterns, i.e. 20 µm grooves on the internal surface and 15 µm rings on the external surface, induced the simultaneous orientation of ECs and SMCs along axial and circumferential directions, respectively, closely mimicking the orthogonally aligned bilayer architecture of natural arteries. Furthermore, the groove patterns significantly accelerated EC migration on the 3D scaffold, potentially promoting endothelial regeneration (or endothelialization) in implanted vascular scaffolds.

## Results

### Working principle of MµCLIP

MµCLIP is a light-induced 3D printing process that uses projected light patterns to induce localized polymerization of the ink (**Figure 1a**). The resolution of the printed structures depends on both lateral (*xy* projection plan) and vertical (*z* axis or printing direction) resolutions. The lateral resolution is primarily governed by the Digital Micromirror device (DMD), which acts as the dynamic mask and has a pixel size of 4.0 µm in our system. The vertical resolution depends on the curing depth *C*_*d*_, which is controlled by the incident light energy and penetration depth *D*_*p*_ of the light into the ink. To achieve a high vertical resolution, the curing depth *C*_*d*_, which is normally set as the layer thickness *L*_*T*_, must be carefully matched to the penetration depth (*D*_*p*_) to prevent the unintentional over-curing or “print-through” (**Figure 1b**). The penetration depth *D*_*p*_ and the optimal incident energy were derived from the Jacob’s working curve^31^, which was obtained by measuring the curing depth *C*_*d*_ as a function of varying UV light exposure (**Supplementary Figure S1**). Using the optimized parameters (*L*_*T*_ = *C*_*d*_ = 80 µm), we attempted to print 20 µm groove patterns. However, the groove patterns were either unresolved or very shallow, measuring only 1-2 µm in depth (**Supplementary Figure S2a**).

**Figure 1.**
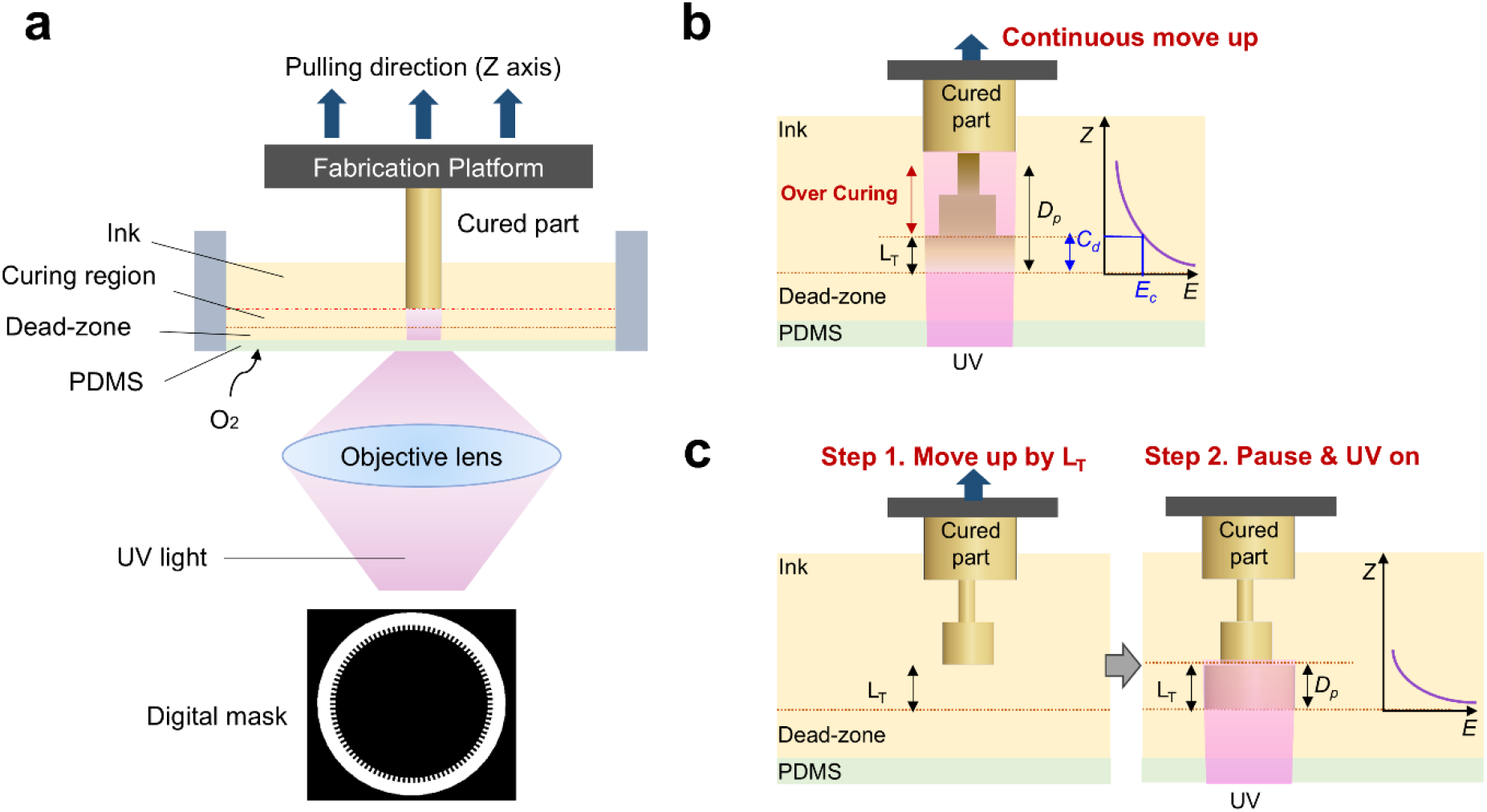
Overview of the setup and working mechanism of the MµCLIP 3D printing method. (**a**) Schematic of the MµCLIP-based 3D printer setup. (**b**) Schematic of the conventional CLIP process, illustrating two key issues: 1) over-curing caused by the mismatch between layer thickness (*L*_*T*_) and penetration depth (*D*_*p*_) of the UV light, and 2) the continuous motion of the fabrication platform during UV projection, which can lead to deformation or loss of fine surface features. (**c**) Schematic of the modified step-wise printing processes with matched *L*_*T*_ and *D*_*p*_ values. This process includes two steps for each layer: the fabrication platform first moves up by the layer thickness (step 1), followed by a pause to allow ink flow before UV projection initiates curing (step 2), ensuring precise pattern resolution.

In the conventional CLIP process, ink curing occurs simultaneously with the continuous upward movement of the fabrication platform, which can induce shear stress and potentially disrupt delicate features, such as surface micropatterns. To mitigate this, we implemented a step-wise printing process. In this approach, the fabrication platform moves upward by the layer thickness (*L*_*T*_) and pauses to allow ink to flow in before initiating UV exposure to cure the current layer (**Figure 1c**). Using this method, we successfully resolved the 20 µm groove patterns, achieving a pattern height of 9.6 µm (**Supplementary Figure S2b**). Our results indicate that both precise matching of *L*_*T*_ to *C*_*d*_ and the step-wise process are crucial for the MµCLIP technique to fabricate large-scale 3D objects with fine topographic micropatterns.

### Fabrication of macroscale scaffolds with microtopographic patterns of various sizes and geometries

To demonstrate the capability of our MµCLIP method to produce microtopographic features, two commonly used pattern geometries, i.e. grooves and pillars, were designed and 3D printed on a simple plate sample using a photocurable citrate-based polymer (CBP) ink. CBP biomaterials were chosen as a model ink for this study due to its biocompatibility, biodegradability, and antioxidant properties as shown in our previous work^32^. Groove patterns, continues along *z*-axis (i.e., the printing direction) were printed to evaluate the lateral resolution (*xy* plane) of the system (**Figure 2a**). We designed groove patterns with varying sizes: width = spacing = 12, 20, 40, and 80 µm, and height = 40 µm, covering the size range of individual mammalian cells (10-50 µm). The printed plate featured four patterned regions and a flat control region. SEM images revealed well-resolved patterns across all regions, closely matching the designed pitch sizes (pitch = width + spacing) (**Supplementary Figure S2e**), which confirmed the high lateral resolution of our MµCLIP process. The printed patterns exhibited a bullet-shape profile with rounded corners, deviating from the designed sharp edges, especially for the smaller patterns (e.g. 12 µm). This is attributed to the Gaussian distribution of the UV light beam, where the intensity diminishes from the beam center to the periphery^31^. Additionally, the pattern heights were lower than the intended 40 µm, ranging from 8.4 µm for the smallest grooves (12 µm width) to 27.8 µm for the largest grooves (80 µm width). This discrepancy is likely due to unintended curing in the recessed regions between patterns caused by light scattering from the surrounding bright areas (i.e., protruding patterns and bulk parts)^33^.

**Figure 2.**
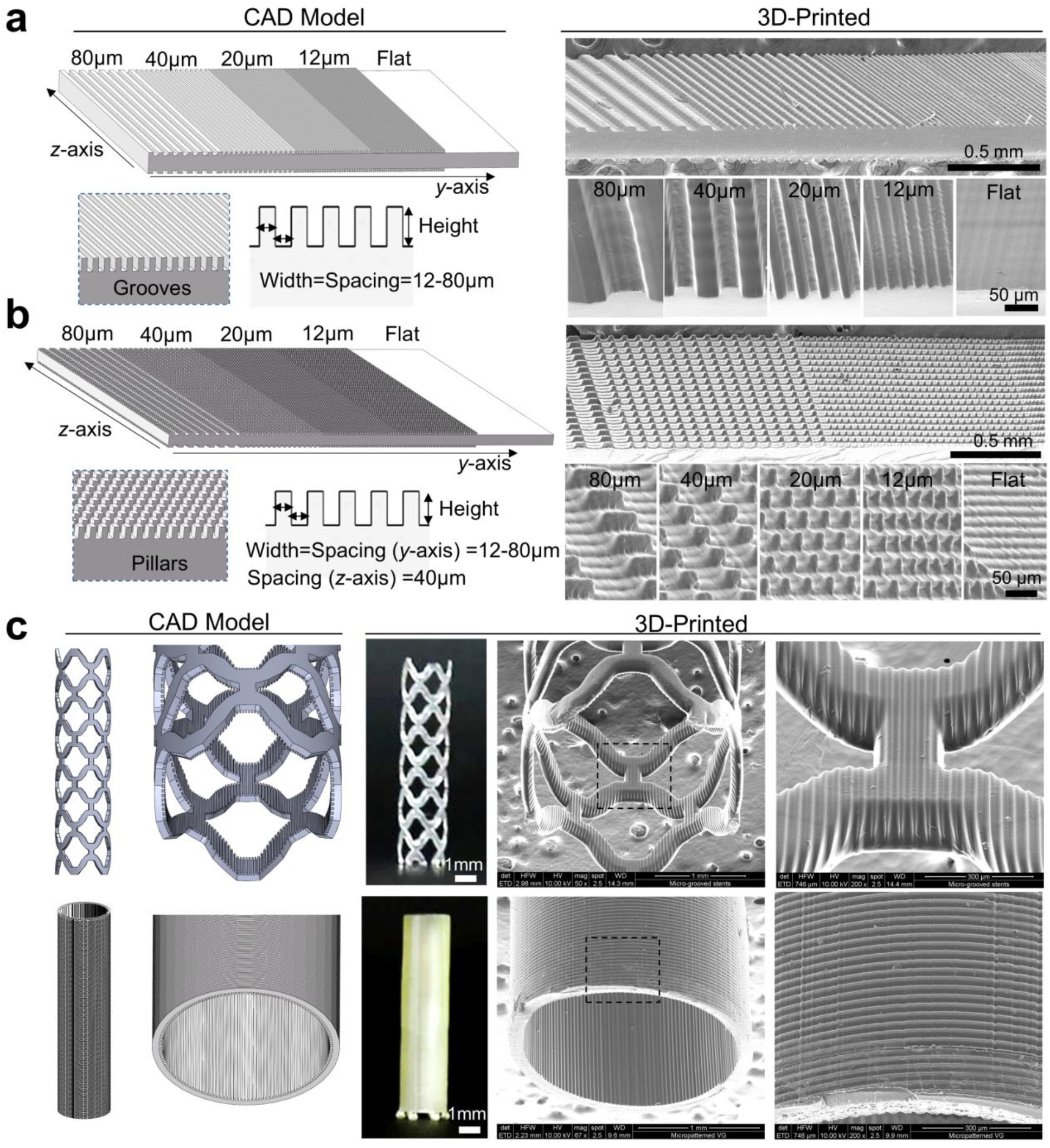
3D-printed plate sample and multi-scale scaffolds with microtopographic patterns of various sizes and geometries. CAD design and scanning electron microscopy (SEM) images of (**a, b**) plate samples with grooves and pillars with varying sizes (12, 20, 40, and 80 µm), and (**c**) multi-scale stent and tubular scaffold (1 cm in length) with hundreds of microns struts (120-130 µm in radial thickness; 250 µm in width) and microtopographic patterns (20 µm grooves; 15 µm rings).

To assess vertical resolution (*z*-axis), pillar patterns with discontinuities along the *z*-axis were printed (**Figure 2b**). Initial prints resulted in pillars that were vertically continuous due to excessive over-curing, with large *D*_*p*_ and *C*_*d*_ values (**Supplementary Figure S2c**). To mitigate this issue, we introduced a light absorber, 2-nitrophenyl phenyl sulfide (NPS), into the CBP ink, which reduced *D*_*p*_ from 80 µm to 33 µm (**Supplementary Figure S1)**. Printing of the ink with reduced *D*_*p*_ produced more discrete vertical structures (**Figure 2b** and **Supplementary Figure S2d**), enabling successful fabrication of pillar patterns with varying widths and spacings (12-80 µm). Similar to the grooves, the heights of the printed pillars were smaller than the designed values.

We further demonstrated the capability of our MµCLIP system by printing multi-scale scaffolds (**Figure 2c**). A stent (1 cm in length) was designed with struts approximately 120 µm thick and 250 µm wide, featuring 20 µm groove patterns on the internal surfaces of the struts. The stent was printed in 10 minutes (1 mm/min). In addition, centimeter-long tubular scaffolds with 130 µm wall thickness were printed, featuring 15 µm ring patterns on the external surface. These results illustrate that our MµCLIP system enables the fabrication of multiscale scaffolds that integrate macroscale architecture, struts in the hundreds-of-microns range, and microtopographic patterns, with hierarchical structures spanning nearly three orders of magnitude in size (i.e., 12 µm to 1 cm).

### Spatial orientation of cells on tubular scaffolds with microtopographic patterns

To investigate the effects of 3D-printed microtopographic patterns on the spatial orientation of cells, tubular scaffolds with 20 µm grooves, 20 µm pillars, or flat internal surfaces (i.e., without patterns) were fabricated (**Figure 3a**). Human umbilical vein endothelial cells (HUVECs) were seeded onto the internal surfaces of these scaffolds. Confocal images revealed that the majority of cells on scaffolds with grooves exhibited a strong alignment along the axial direction of the scaffold (**Figure 3b**). In contrast, cells on scaffolds with pillar patterns or flat surfaces showed random orientations, as confirmed by quantification through polar diagrams. This preferential alignment on grooved surfaces may support normal endothelial functions, closely replicating the structural organization of the endothelium in native arteries. Immunostaining for CD144 (VE-cadherin), a key marker for endothelial cell junctions, showed that cells on scaffolds with grooves displayed well-organized and strong junctional expression among aligned cells (**Figure 3c**). Conversely, cells on scaffolds with pillars or flat surfaces exhibited more disorganized CD144 structures, indicative of less structured cell junctions.

**Figure 3.**
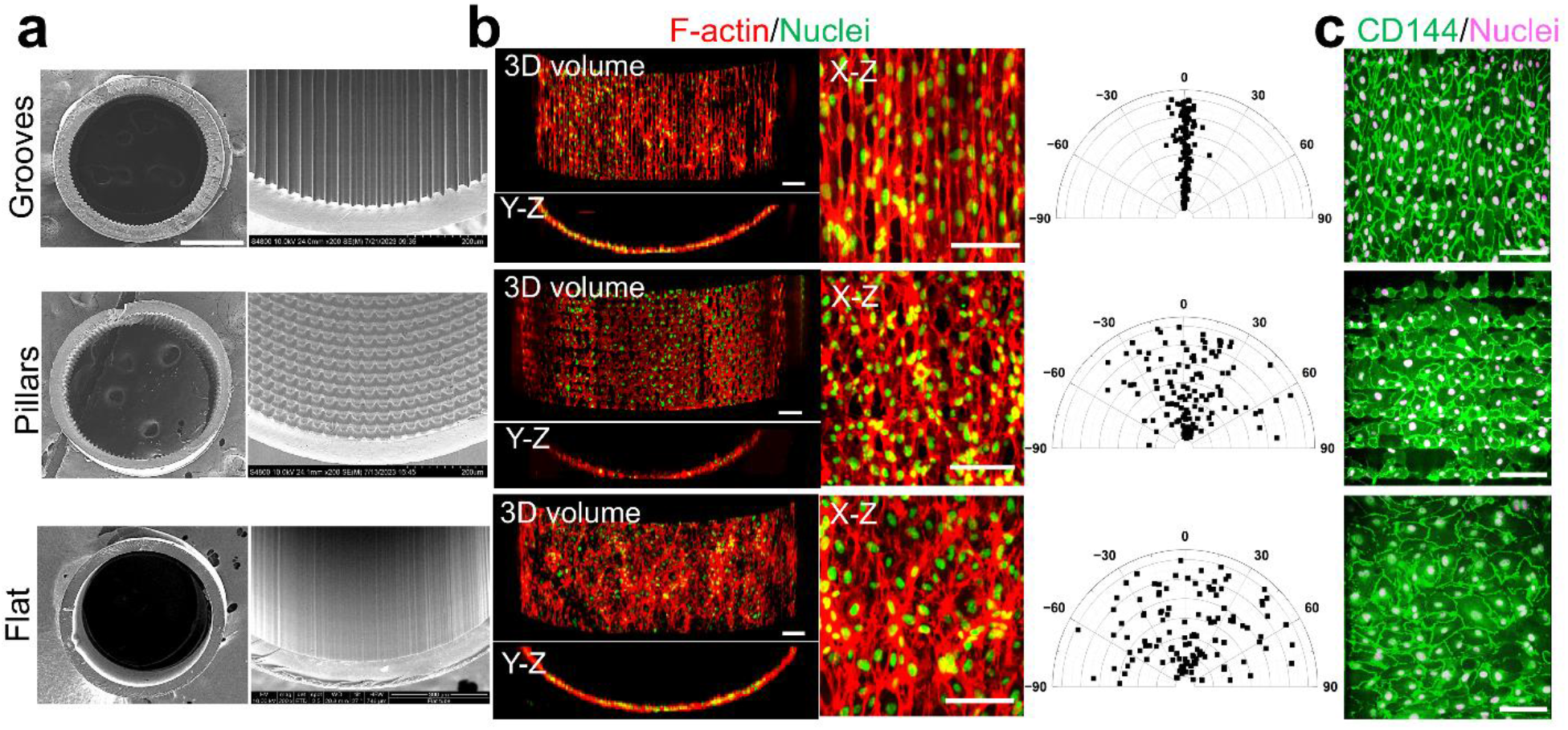
3D-printed scaffolds with microtopographic patterns guide distinct tissue organization. (**a**) SEM of tubular scaffolds with 20 µm grooves, 20 µm pillars, or flat surfaces on internal surfaces. (**b**) Confocal fluorescent images showing the orientation of endothelial cells (HUVECs) seeded on the internal surfaces of 3D-printed scaffolds. The polar diagrams quantify cell orientation, with over 120 cells measured from n = 4 images for each type of scaffold. (**c**) Immunostaining of functional marker, e.g. CD144 (or VE-cadherin), of cells on scaffolds. Scale bars: 200 μm.

### Concurrent, distinct orientation of multiple cell types in a single scaffold with dual patterns

Achieving concurrent control over the spatial organization of multiple cell types within a single scaffold is crucial for regenerating or repairing complex tissues and organs; however, it remains an enduring challenge. Our 3D MµCLIP system enables the simultaneous fabrication of multiple types of microtopographic patterns on a single scaffold. As a proof of concept, we designed a tubular scaffold with axially aligned 20 µm grooves on the internal surface and circumferentially aligned 15 µm rings on the external surface to replicate the orthogonally aligned bilayer architectures of the natural arterial wall (**Figure 4a**). SEM images showed the successful fabrication of both groove patterns on the internal surface and ring patterns on the external surface of the scaffold (**Figure 4a**). After seeding HUVECs on the internal surface and human aortic smooth muscle cells (HASMCs) on the external surface of the scaffold, the micropatterns induced strong axial alignment of HUVECs and circumferential alignment of HASMCs (**Figure 4b**), closely mimicking the structures of the intima and media layers in natural arteries. In contrast, HUVECs on internal surfaces of flat scaffolds showed random organization while HASMCs exhibited spontaneous alignment along axial direction. In addition, HUVECs grown on micropatterned and flat scaffolds exhibited the formation of mature cell-cell junctions (**Figure 4c**). While HASMCs grown on both scaffolds expressed high levels of α-smooth muscle action (α-SMA), the cells on the scaffolds with ring patterns demonstrated more densely packed and organized α-SMA structures.

**Figure 4.**
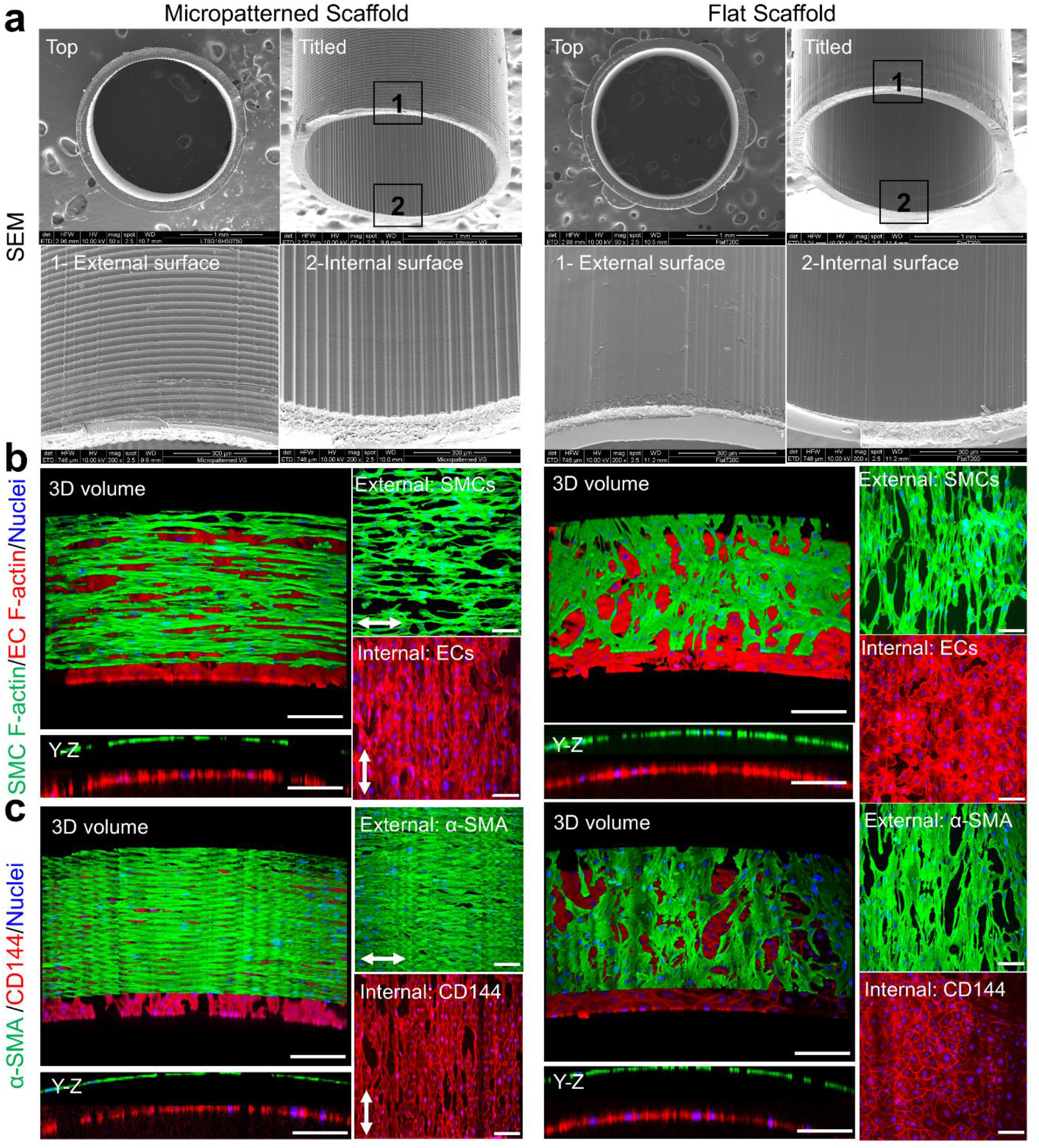
3D-printed scaffolds with dual patterns guide simultaneous orientation of ECs and SMCs along axial and circumferential directions, respectively. (**a**) SEM images of 3D-printed scaffolds with dual patterns (i.e., 15 µm rings on the external surface and 20 µm grooves on the internal surface) and flat surfaces as the control. **(b)** Confocal fluorescent images displaying F-actin and nuclei of ECs and SMCs grown on internal and external surfaces of the scaffolds, respectively. **(c)** Confocal fluorescent images showing CD144 (or VE-Cadherin) of ECs and α-SMA of SMCs grown on the internal and external surfaces of the scaffolds, respectively. Scare bars in **b** and **c** are 100 µm. Arrows indicate directions of micropattern alignment and corresponding cell orientation.

### Accelerated cell migration on 3D tubular scaffold with groove patterns

Migration of host cells into implanted scaffolds is essential for rapid repopulation and in-situ tissue regeneration^34, 35^. For instance, evidence indicates that migration of host ECs or endothelial progenitor cells from the injury or anastomotic sites significantly contributes to endothelial regeneration (or endothelialization) on vascular grafts^36-38^. Although chemical modifications, such as immobilization of growth factors or peptides, have been employed to promote endothelialization on vascular grafts, incomplete endothelization, particularly in the central region of vascular grafts, remains a persistent issue^39-41^. Controlling cell migration in 3D-structured scaffolds is particularly challenging. To investigate the effect of microtopographic patterns on cell migration within a 3D environment, we fabricated a tubular scaffold with half of its internal surface featuring a grooved region and the other half a flat region (**Figure 5a**). HUVECs were seeded at one end of the scaffold, and their proliferation and migration towards the opposite end were monitored over time (**Figure 5b**). Initially, cells were evenly distributed across both the grooved and flat regions. Cells on the grooved surfaces exhibited strong alignment and elongation along the groove direction, while cells on the flat regions were randomly spread. By day 1 and day 3, the leading edge of cells on the grooved region had migrated slightly further than cells on the flat region. By day 6, the difference became more pronounced, with cells on the grooved region advancing approximately 1700 µm ahead of those on the flat region (**Figure 5b, 5c**). Additionally, cells on the grooved surface remained elongated and aligned throughout the 6-day culture period. These results demonstrate that groove patterns can significantly accelerate cell migration along the groove direction on a 3D tubular scaffold. This suggests that microtopographic patterns may facilitate the rapid repopulation of implanted scaffolds through enhanced host cell migration, supporting in-situ tissue regeneration, such as the rapid endothelialization of vascular scaffolds.

**Figure 5.**
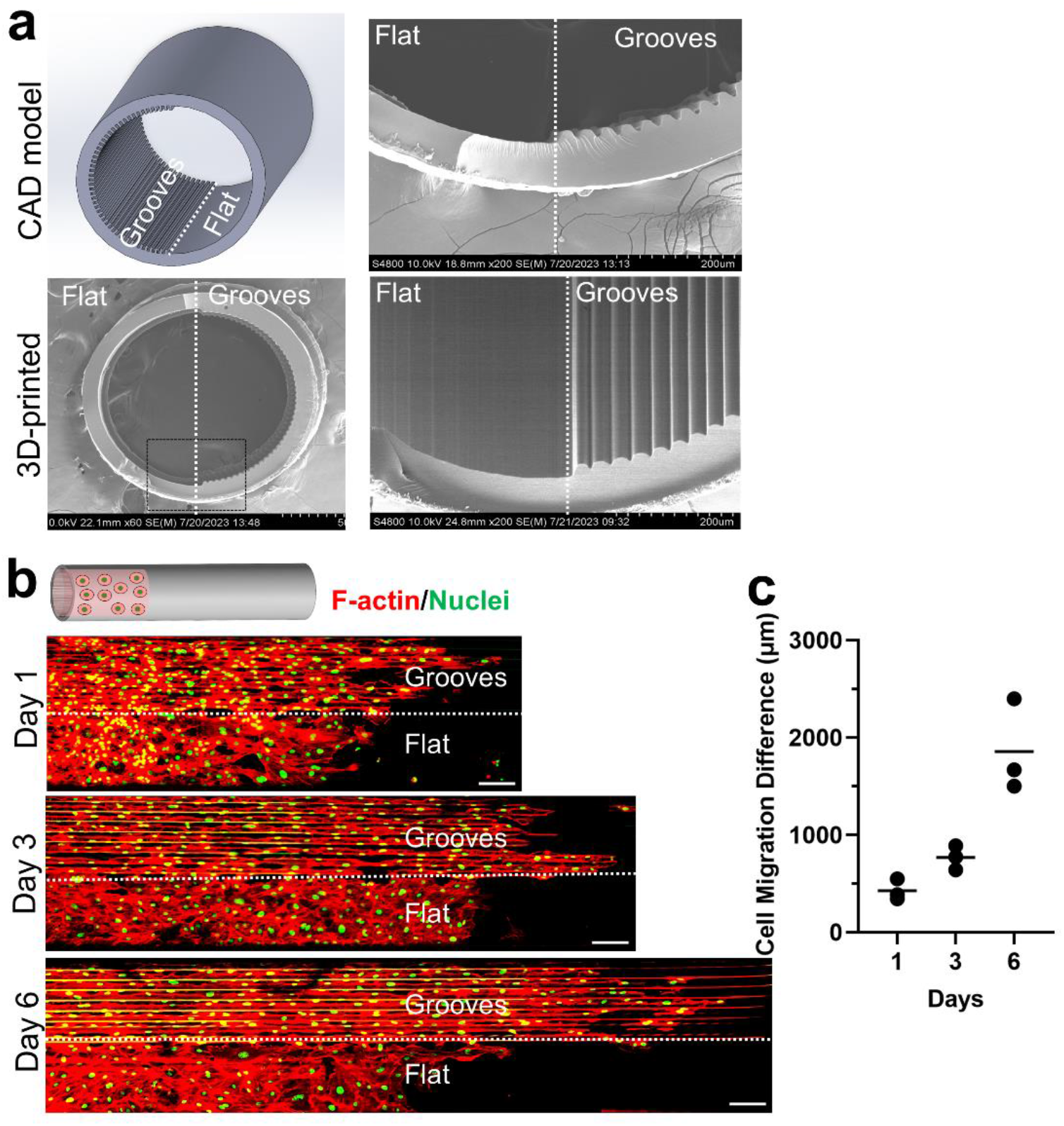
3D-printed tubular scaffolds with 20 µm groove patterns accelerated EC migration. **(a)** CAD model and SEM images of 3D-printed tubular scaffolds with internal surfaces divided into patterned and flat regions. The two rightmost SEM images are magnified views of the scaffold (black box). **(b)** Confocal fluorescence images showing EC migration on internal scaffold surfaces over a 6-day period. The schematic illustrates the seeding of cells at one end of the scaffold on day 0. Scale bars: 200 μm. **(c)** Quantification of the difference in migration distance, measured as the distance between the leading edges of cells on the grooved and flat surfaces. n = 3, with lines indicating mean values.

## Discussion

In this study, we introduced a high-speed, multiscale 3D printing method - MµCLIP to enable the rapid fabrication of centimeter-scale 3D scaffolds featuring microtopographic patterns. Our method offers several advantages over existing approaches. First, MµCLIP provides precise control over microtopographic patterns in terms of geometry, size, and spatial location within macroscale 3D structures. While phase separation techniques have been used to create nanostructures within 3D-printed objects, they offer limited control over shape (e.g., irregularly shaped pores) and spatial distribution (e.g., random distribution throughout the object)^25, 26^. In contrast, our 3D printing method allows the creation of highly ordered arrays of microtopographic patterns with a wide range of geometries (e.g., grooves, pillars, and rings) and sizes (e.g., 12–80 µm). Moreover, distinct patterns can be strategically positioned on specific areas of a single scaffold. This capability is essential for designing scaffolds with highly controlled 3D microenvironments that guide complex organization of cells and tissues. Second, MµCLIP can fabricate hierarchical structures with feature sizes spanning three orders of magnitude (12 µm-1 cm), which was rarely achieved^28, 29, 42, 43^. As shown in Figure 2, we produced scaffolds measuring one centimeter in length, with struts on the scale of hundreds of microns and microtopographic patterns. This capacity to generate hierarchical structures expands the range of potential applications, particularly for implants that need to mimic the multiscale architecture of natural tissues, such as bone or vascular systems. Third, the speed of MµCLIP (1 mm/min) significantly outpaces other multiscale fabrication techniques. The trade-off between high resolution and printing speed has been the long-standing challenge in multiscale fabrication. For instance, many extrusion-based multiscale printers are constrained by slow print speeds due to the long print paths^28, 29^. In comparison, DeSimone’s group demonstrated a CLIP-based high-resolution 3D printer that printed a 2-cm-long lattice structure with 100 µm-thick struts in 1.5 hours, i.e. 0.2 mm/min, which is 10^5^ times faster than two-photon-based Nanoscribe printer^30^. Notably, our printing speed is five times faster than theirs. Together, our MµCLIP method provides an unparalleled combination of speed, precision, and versatility in fabricating multiscale hierarchical structures.

The multiscale biomaterial scaffolds produced by our MµCLIP method hold significant potential for tissue repair or regeneration by offering distinct functional advantages across multiple scales: 1) anatomically accurate overall shape (millimeters to centimeters) for perfect matching and integration with biological system, 2) 3D-structured struts in hundreds of microns scale for providing necessary mechanical support, and 3) microtopographic patterns (submicron to tens of microns) for guiding cell behavior (e.g., alignment) and tissue function. To showcase the potential of this approach, we fabricated tubular scaffolds with multiscale structures and demonstrated their applicability for biomimetic arterial regeneration.

In vascular tissue engineering, recapitulating the multi-layered, orthogonally orientated structures of natural blood vessels has emerged as a promising strategy for developing biomimetic vascular grafts and more accurate *in vitro* models of arteries. While the ECs on the inner layer of vessels are naturally aligned axially due to the blood flow shear stress, much less is understood about how to achieve circumferential alignment of SMCs in the medial layer^44, 45^. Engineering strategies to induce circumferential alignment of SMCs remain limited. Therefore, simultaneously achieving orthogonal alignment of ECs and SMCs has been a long-standing challenge. Some studies have utilized sheet-based methods to grow individually aligned cell sheets, which are then rolled into a multi-layered tube^46, 47^. Another approach employed freezing-indued alignment of electrospun fibers to promote axial alignment of ECs, while encapsulating SMCs within a hydrogel layer to achieve spontaneous circumferential alignment^18^. However, these existing methods often involve lengthy, complicated, multi-step fabrication processes. By contrast, our MµCLIP method enables the rapid, one-step fabrication of bioresorbable scaffolds with dual microtopographic patterns that induce simultaneous alignment of ECs and SMCs along axial and circumferential directions, respectively, closely replicating the native architecture of arteries. Additionally, the acceleration of cell migration on scaffolds with groove patterns could enhance the repopulation of implanted scaffolds by host cells, e.g. facilitating rapid endothelization, which is a critical yet challenging requirement for blood-contacting implants. These 3D-printed scaffolds with anisotropic microtopographic patterns provide robust guidance for the spatial orientation of cells and tissues within a 3D-structured environment. This capability not only advances the biomimetic regeneration of arteries but also holds promise for regenerating other oriented tissues, such as nerves, muscles, myocardium and intestines.

In addition to the 3D printing of anisotropic microtopography for guiding cellular and tissue orientation, we also demonstrated the 3D printing of isotropic pillar patterns (Figure 2b). Recent work by Wang et al. showed that microtopographic pillars can induce nuclear deformation in human mesenchymal stem cells, promoting osteogenic differentiation and accelerating bone regeneration in a rodent cranial defect model^14^. However, their patterned implants were restricted to 2D planar films due to the limitations of the lithography-based manufacturing method used to create micropillars. Similarly, another study reported that silicone breast implants with topographic features (random structures with average roughness of 4 µm) significantly reduced the inflammation and foreign body responses in mouse and rabbit models compared to implants with smooth surfaces or larger topographic features^48^. These findings underscore the importance of microtopography in scaffolds and medical devices, suggesting they can be strategically engineered not only to guide cellular and tissue alignment but also to influence broader tissue responses and integration *in vivo*. By overcoming the challenge of creating microtopography on curved 3D scaffold surfaces, our approach provides a powerful tool for designing advanced scaffolds and medical devices that harness microtopography to enhance therapeutic outcomes.

## Methods

### Polymer synthesis and ink preparation

Photocurable citrate-based polymer (CBP) was synthesized and characterized by following our previously reported protocol^32^. Briefly, citric acid and 1,12-dodecanediol at molar ratio of 2:3 were melted (165 °C, 22 min), co-polymerized (140 °C, 60 min), purified, and freeze-dried to yield poly (1,12-dodecanediol citrate) (PDC) pre-polymer. Every 22 g PDC pre-polymer was dissolved in tetrahydrofuran (180 mL) with imidazole (816 mg) and glycidyl methacrylate (17.8 mL), reacted (60 °C, 6 h), purified, and freeze-dried to yield methacrylate PDC (mPDC) pre-polymer. The characterization of mPDC pre-polymer was performed by proton nuclear magnetic resonance (^1^H NMR) using deuterated dimethyl sulfoxide (DMSO-*d*_6_) as the carrier solvent.

To formulate CBP base ink for 3D printing, 75 wt% mPDC pre-polymer was mixed with 2.2 wt% Irgacure 819, acting as a primary photoinitiator, and 3.0 wt% Ethyl 4-dimethylamino benzoate (EDAB), acting as a co-photoinitiator, in a solvent of pure ethanol. The CBP based inks were used to print scaffolds without micropatterns or with groove patterns. To print scaffolds with pillar or ring patterns, the 0.5 wt% NPS was introduced as the light absorber and mixed with the base ink for the control of penetration depth *D*_*p*_ of the light into the ink.

### Working curve measurement of the ink

The 3D printing parameters for different microtopographic patterns were obtained through experimental measurements of working curve equations (Eq. (1)) for inks.

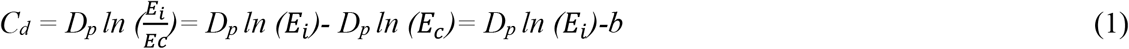

Where *C*_*d*_ is Curing depth, *Ec* corresponds to the threshold of the transition from liquid to solid upon polymerization, *D*_*p*_ is penetration depth, equals to the reciprocal of attenuation coefficient, b is the intercept of *C*_*d*_.

Next, the systematic energy to print each layer was computed based on working curve equation. After arrangement of Eq. (1), E_i_ can be obtained as follows,

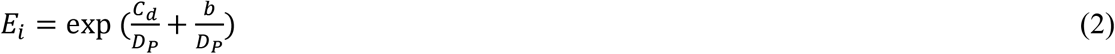

To compensate the energy decay of the light through dead zone (*DZ*), the systematic incident energy *E*_0_ should be described as Eq. (3),

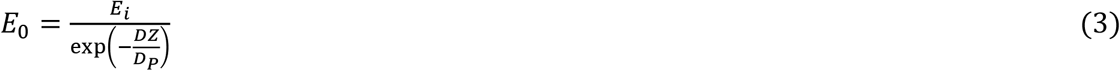

The *DZ* thickness is approximated as 40 µm based on our measurement following previously published method^24^.

To measure the working curves, a cover glass was placed on oxygen permeable window made of polydimethylsiloxane (PDMS) in the resin tank, then a droplet of ink was put on the top of the cover glass. That was followed by projecting UV light with varying dosage, in which dosage is the product of power density and exposure time. The power density is constant while the exposure time is varied. Next, the cured layer on the glass was rinsed by pure ethanol to remove uncured resin. The cured thickness was measured by micrometers. At least four cured thicknesses for inks at various dosages were measured to plot working curves. The working curve equations were obtained from the logarithmic trendline of data points in excel.

### Fabrication and characterization of scaffolds with microtopographic patterns

3D printing was performed using a homemade micro-continuous liquid production process (µCLIP)-based printer. An oxygen-permeable window made of PDMS was attached to the bottom of the resin vat. A digital micromirror device (DMD, Texas Instruments Inc., Plano, TX) was utilized as the dynamic mask generator to pattern the UV light (365 nm). Projection optics of the printer were optimized to have a pixel resolution of 4.0 μm × 4.0 μm at the focal plane. CAD files of the print part were designed in SolidWorks (Dassault Systèmes, Waltham, MA). The resulting STL files were sliced for a 2D file output by CHITUBOX 3D slicer software. Full-screen images were projected onto the vat window for photopolymerization. The base cure time was 20 s for the initial attachment of 3D-printed parts to the fabrication platform. The power intensity and exposure time for each layer in terms of different types of patterns were listed in **Supplementary Table S1**. Between layers, pause time up to 0.8 s with solid black image projected may be implemented to allow replenishment of ink to in the dead zone depending on the viscosity of the ink. After printing, the parts were rinsed in ethanol to remove the unpolymerized ink and moved into a heating oven for thermal curing at 80 °C for 24 hours.

The structure and surface morphology of the BVSs were observed by scanning electron microscope (SEM; Hitachi S4800, Japan). Strut and micropattern sizes were measured from SEM images.

### Cell Culture

Human umbilical vein endothelial cells (HUVECs; ATCC, Manassas, VA) were expanded in growth media consisting of Endothelial Cell Growth Kit-VEGF (ATCC PCS-100-041) in Vascular Cell Basal Medium (ATCC PCS-100-030) under the standard culture condition (37 °C with 5 % CO2 in a humid environment) to 80 % confluency before passaging. HUVECs at passages 5–7 were used. Human aortic smooth muscle cells (HASMCs; Cell Applications, Inc., San Diego, CA) were expanded in human SMC growth medium kit (Cell Applications, Inc., 311K-500) under the standard culture condition (37 °C with 5 % CO2 in a humid environment) to 80 % confluency before passaging.

### Cell seeding, immunostaining and imaging

For the culture of HUVECs on the internal surfaces of 3D-printed tubular scaffolds, 8 µL of suspended HUVECs (200 × 10^4^ cells/ml) were injected to fill the entire lumen of the tubular scaffolds. After a 5-minute incubation for cell attachment, the scaffold was gently rotated by 90°, followed by a 30-minute incubation. The cell suspension was then carefully aspirated from the scaffold. After rotating the scaffold another 90°, another 8 µL of suspended HUVECs (200 × 10^4^ cells/ml) was injected into the lumen of the tubular scaffold, and the same seeding process was repeated. Finally, the tubular scaffolds with attached cells were transferred into pre-warmed growth medium.

For the co-culture of HUVECs and HASMCs on the 3D-printed tubular scaffolds with dual microtopographic patterns, a custom, 3D-printed co-culture device with a double-tube structure was used to simultaneously seed both cell types (**Supplementary Figure S3**). The inner tube was the scaffold with or without microtopographic patterns. The scaffold was connected to an outer tube (inner diameter: 4 mm, wall thickness: 200 μm) via cuboid supports sizing 250 μm, spaced 950 μm apart. The HASMC suspension (30 × 10^4^ cells/ml) was seeded into the space between the inner and the outer tubes, while the HUVEC suspension (20 × 10^4^ cells/ml) was seeded into the inner tube. To ensure uniform cell distribution on scaffolds, the co-culture platform was rotated 45 degrees clockwise at intervals of 7–10 minutes upon seeding. Following this, the platform, containing both HASMCs and HUVECs, was transferred to a co-culture medium composed of equal parts HASMC and HUVEC growth medium. The cells were cultured for 36 hours to characterize F-actin, and for 3 days to evaluate the expression of functional markers.

Cells on scaffolds were fixed with 4 % formaldehyde (Fisher Scientific, Fair Lawn, NJ) for 10 min and permeated with 0.1 % Triton X-100 for 15 min. To examine the cell morphology, the cells were incubated with Alexa Fluor 594 Phalloidin (10 μM; A12381; ThermoFisher Scientific, Fair Lawn, NJ) and SYTOX Green Nucleic Acid Stain (0.15 μM; S7020; ThermoFisher Scientific) in PBS solution with 1.5 % bovine serum albumin (BSA) for 30 min. To examine the cell phenotype, cells were firstly incubated with 1.5 % BSA for 40 minutes to block non-specific background following fixation and permeation. Next, HASMCs were incubated in sequence with Recombinant Anti-alpha smooth muscle Actin primary antibody (0.572 μg/ml; ab133567; Abcam Inc., MA) (α-SMA) and HUVECs were incubated with Anti-VE Cadherin (or CD144) primary antibody (3.33 μg/ml; ab33168; Abcam Inc.) overnight at 4 °C, then both cells were incubated with SYTOX Green Nucleic Acid Stain (0.15 μM; S7020; ThermoFisher Scientific) and Alexa Fluor 594 Donkey anti-Rabbit IgG (H+L) secondary antibody (4 μg/ml; A21207; ThermoFisher Scientific) in PBS solution with 1.5 % BSA for 60 min at room temperature. The samples were finally washed three times in PBS. Prior to confocal imaging, the outer tube and supports were removed by micro-scissors and the inner tubes were placed on microscope slides for examination using Nikon spinning disk confocal microscope.

### Cell migration

HUVECs were seeded into one end of the tubular scaffolds (∼ 1/3 of the total length), with half of the internal surface featuring groove patterns and the other half a flat region. Following the same seeding protocol described in the previous section, 3 µL of the suspended HUVECs (200 × 10^4^ cells/ml) were injected twice into the scaffold lumen. After seeding, the HUVECs were cultured in the growth medium. After culturing for 1, 3 and 6 days, the cells were fixed, permeated, stained with the Alexa Fluor 594 Phalloidin and SYTOX Green Nucleic Acid Stain, and imaged using Nikon spinning disk confocal microscope.

### Statistics and reproducibility

Unless otherwise specified, data was presented as mean ± standard deviation (s.d.). For each experiment, at least three samples were analyzed. One-way ANOVA with Tukey’s Multiple Comparison Test was used to analyze statistical significance. A *P* value of <0.05 was considered to indicate a statistically significant difference. All experiments presented in the manuscript were repeated at least as three independent experiments with replicates to confirm the results are reproducible.

## Acknowledgements

This work was supported by American Heart Association Career Development Award (AHA, Grant: 852772). The authors gratefully acknowledge Pengpeng Zhang for his technical support in MATLAB to generate digital masks for 3D printing.

